# Comparison of single genome and allele frequency data reveals discordant demographic histories

**DOI:** 10.1101/182899

**Authors:** Annabel C. Beichman, Tanya N. Phung, Kirk E. Lohmueller

**Author notes:** **To whom correspondence should be addressed:** Kirk E. Lohmueller, Department of Ecology and Evolutionary Biology, University of California, Los Angeles, 621 Charles E. Young Drive South, Los Angeles, CA 90095-1606.

## Abstract

Inference of demographic history from genetic data is a primary goal of population genetics of model and non-model organisms. Whole genome-based approaches such as the Pairwise/Multiple Sequentially Markovian Coalescent (PSMC/MSMC) methods use genomic data from one to four individuals to infer the demographic history of an entire population, while site frequency spectrum (SFS)-based methods use the distribution of allele frequencies in a sample to reconstruct the same historical events. Although both methods are extensively used in empirical studies and perform well on data simulated under simple models, there have been only limited comparisons of them in more complex and realistic settings. Here we use published demographic models based on data from three human populations (Yoruba (YRI), descendants of northwest-Europeans (CEU), and Han Chinese (CHB)) as an empirical test case to study the behavior of both inference procedures. We find that several of the demographic histories inferred by the whole genome-based methods do not predict the genome-wide distribution of heterozygosity nor do they predict the empirical SFS. However, using simulated data, we also find that the whole genome methods can reconstruct the complex demographic models inferred by SFS-based methods, suggesting that the discordant patterns of genetic variation are not attributable to a lack of statistical power, but may reflect unmodeled complexities in the underlying demography. More generally, our findings indicate that demographic inference from a small number of genomes, routine in genomic studies of nonmodel organisms, should be interpreted cautiously, as these models cannot recapitulate other summaries of the data.

## INTRODUCTION

The Pairwise Sequentially Markovian Coalescent (PSMC) and related methods have become a popular tool to estimate the history of a population from genetic variation data (McVean and Cardin 2005; Li and Durbin 2011; Schiffels and Durbin 2014). These methods use whole genome sequences from one to four individuals to infer the demographic history of an entire population. Specifically, they estimate the local time to the most recent common ancestor (TMRCA) for small regions in the genome, then use the distribution of these coalescent times to infer an overarching demographic history. For instance, if many regions of the genome coalesce at a specific time, it may be evidence for a population contraction, which would reduce the number of genetic lineages. The great appeal of these methods is that they do not rely on deep sequencing of multiple individuals in a population; instead, a single genome can be used to infer the demographic history of an entire population. PSMC and its successors have been used to infer the demographic histories and split times of many human populations (Li and Durbin 2011; Kidd *et al.* 2012; Schiffels and Durbin 2014; 1000 Genomes Project Consortium 2015; Henn *et al.* 2016), and were recently featured in three prominent articles that reconstructed human history using whole genome sequencing data from over 20 populations (Malaspinas *et al.* 2016; Mallick *et al.* 2016; Pagani *et al.* 2016).

PSMC plots have also become a cornerstone of many studies of non-model organisms lacking resources for the sequencing of numerous individuals, including archaic hominins (Meyer *et al.* 2012; Prufer *et al.* 2014), great apes (Prado-Martinez *et al.* 2013), wild boars and domestic pigs (Groenen *et al.* 2012; Bosse *et al.* 2014), canids (Freedman *et al.* 2014; Wang *et al.* 2016), horses (Orlando *et al.* 2013), over 38 bird species (Nadachowska-Brzyska *et al.* 2013; Hung *et al.* 2014; Nadachowska-Brzyska *et al.* 2015; 2016; Murray *et al.* 2017), pandas (Zhao *et al.* 2012), dromedaries (Fitak *et al.* 2016), flowering plants (Albert *et al.* 2013; Ibarra-Laclette *et al.* 2013; Holliday *et al.* 2016), and even woolly mammoths (Palkopoulou *et al.* 2015).

Despite their wide-spread prominence, there is concern over the validity of demographic models obtained from this set of whole genome-based methods. Particularly, Mazet *et al.* (2015) found that PSMC captures the inverse instantaneous coalescent rate (IICR) rather than an absolute measure of population size. The IICR corresponds to the effective population size if the population is panmictic, but it can differ from the population size due to gene flow and population structure which affect the time to coalescence between subgroups. Thus, population structure can give a false signal of population growth or contraction - a notorious problem in demographic inference (Ptak and Przeworski 2002; Chikhi *et al.* 2010; Peter *et al.* 2010; Gattepaille *et al.* 2013; Heller *et al.* 2013; Mazet, Rodriguez, and Chikhi 2015; Mazet, Rodriguez, Grusea, *et al.* 2015; Orozco-terWengel 2016). Given these possible confounders, the degree to which whole genome-based plots derived from PSMC and its successors correspond to actual population size changes, rather than other demographic phenomena, remains unclear.

An alternative approach to infer population demography from genetic data uses the site frequency spectrum (SFS). The SFS represents the distribution of alleles at different frequencies in a sample of individuals from a population (Nielsen 2000; Wakeley 2009). The distribution of single nucleotide polymorphisms (SNPs), ranging from rare ‘singletons’ which appear only once in the sample, to high-frequency variants that may appear in the majority of individuals, is directly affected by the demographic history of the population (Nielsen 2000; Wakeley 2009). Population contractions (‘bottlenecks’) can lead to a dearth of rare variants (Nei *et al.* 1975), whereas a rapid population expansion can lead to an overabundance (Tajima 1989; Slatkin and Hudson 1991; Keinan and Clark 2012). The SFS is a sufficient statistic for unlinked SNPs and has been used extensively in population genetic inference of demography (Nielsen 2000; Polanski and Kimmel 2003; Adams and Hudson 2004; Marth *et al.* 2004; Keinan *et al.* 2007; Gutenkunst *et al.* 2009; Gravel *et al.* 2011; Excoffier *et al.* 2013). SFS-based demographic inference has been implemented in programs such as *dadi* (Gutenkunst *et al.* 2009), moments (Jouganous *et al.* 2017), fastsimcoal2 (Excoffier *et al.* 2013), stairway plot (Liu and Fu 2015), fastNeutrino (Bhaskar *et al.* 2015), and others (Schraiber and Akey 2015). The SFS requires less sequence data per individual than the whole genome methods, but requires a greater number of individuals to be studied, with a minimum of ten per population typically used (Gutenkunst *et al.* 2009; Excoffier *et al.* 2013). While the SFS is impractical if one can only sequence one or two individuals per population, population genomic studies based on many short loci scattered throughout the genome are beginning to be carried out on non-model organisms. RAD-seq data or gene transcript data from RNA-seq can readily be used for SFS-based demographic inference (McCoy *et al.* 2014; Trucchi *et al.* 2014; Sovic *et al.* 2016).

SFS-based and whole genome-based methods may have different strengths and weaknesses for demographic inference (Schraiber and Akey 2015). Theoretical and empirical data show that SFS-based approaches using large numbers of individuals can accurately estimate recent population growth (Nelson *et al.* 2012; Tennessen *et al.* 2012; Gazave *et al.* 2013; Bhaskar *et al.* 2015; Gao and Keinan 2016). In contrast, whole genome-based methods are less able to do so (Li and Durbin 2011). Recently, however, Schiffels and Durbin (2014) developed the multiple sequentially Markovian coalescent (MSMC), an extension to PSMC that uses the SMC’ algorithm (Marjoram and Wall 2006) and can infer demography from two, four or eight haplotypes (also known as PSMC’ when inferring from two haplotypes). The incorporation of multiple genomes in MSMC is specifically meant to improve estimates of recent growth (Schiffels and Durbin 2014).

The SFS may be limited in the degree to which it can detect ancient bottlenecks > *2Ne* (effective population size) generations ago and in its ability to detect population declines (Bunnefeld *et al.* 2015; Terhorst and Song 2015; Boitard *et al.* 2016). Whole genome-based approaches are not constrained *a priori* by the number of population size changes as is common in the SFS-based approaches (but see the “stairway plot” approach of Liu and Fu (2015)). They therefore often give information about events occurring millions of years ago, but the reliability of those results remains uncertain (Li and Durbin 2011). Further, demographic models inferred from human populations using the SFS were unable to recapitulate the empirical distribution of identity by state (IBS) tracts across the genome, while PSMC-derived models and a new IBS-derived model were better able to match the IBS tract distribution (Harris and Nielsen 2013). However, the IBS-derived model did not predict the empirical SFS.

Due to these different strengths and weaknesses of approaches using a single type of data, new methods have been developed which attempt to combine linkage disequilibrium (LD) information and the SFS (Bunnefeld *et al.* 2015; Boitard *et al.* 2016; Terhorst *et al.* 2017; Weissman and Hallatschek 2017). One of the most recent is Terhosrt, Kamm and Song’s (2017) method, SMC++, which combines a PSMC-like approach with the SFS to condition an SFS calculated from many individuals on the distribution of TMRCA from a single unphased genome. This approach is fast and potentially very powerful, but has the same barrier to entry for those studying non-model organisms as the other SFS methods, as it requires sequence data from many individuals.

Due to anthropological and biomedical interest, humans are an organism that has been extensively studied using numerous demographic inference methods and provide a means to quantitatively compare these demographic inference approaches using the same empirical populations. Gutenkunst *et al.* (2009) and Gravel *et al.* (2011) carried out SFS-based inference of human demography using the diffusion approximation in ∂a∂i, while Li and Durbin (2011) and Schiffels and Durbin (2014) estimated human demography from the same populations using PSMC and MSMC, respectively. Although the results are in some ways generally similar, the demographic models inferred for three human populations using MSMC (Schiffels and Durbin 2014) differ from demographic models for the same populations derived from SFS-based methods (Gutenkunst *et al.* 2009). MSMC infers ancient ancestral sizes and periods of growth and decline (the characteristic “humps” in MSMC trajectories) that were not detected in the SFS-derived models as well as inferring greater recent growth (Figure 1). The models inferred using MSMC also vary depending on the number of genomes used for the inference (Figure 1).

**Figure 1.**
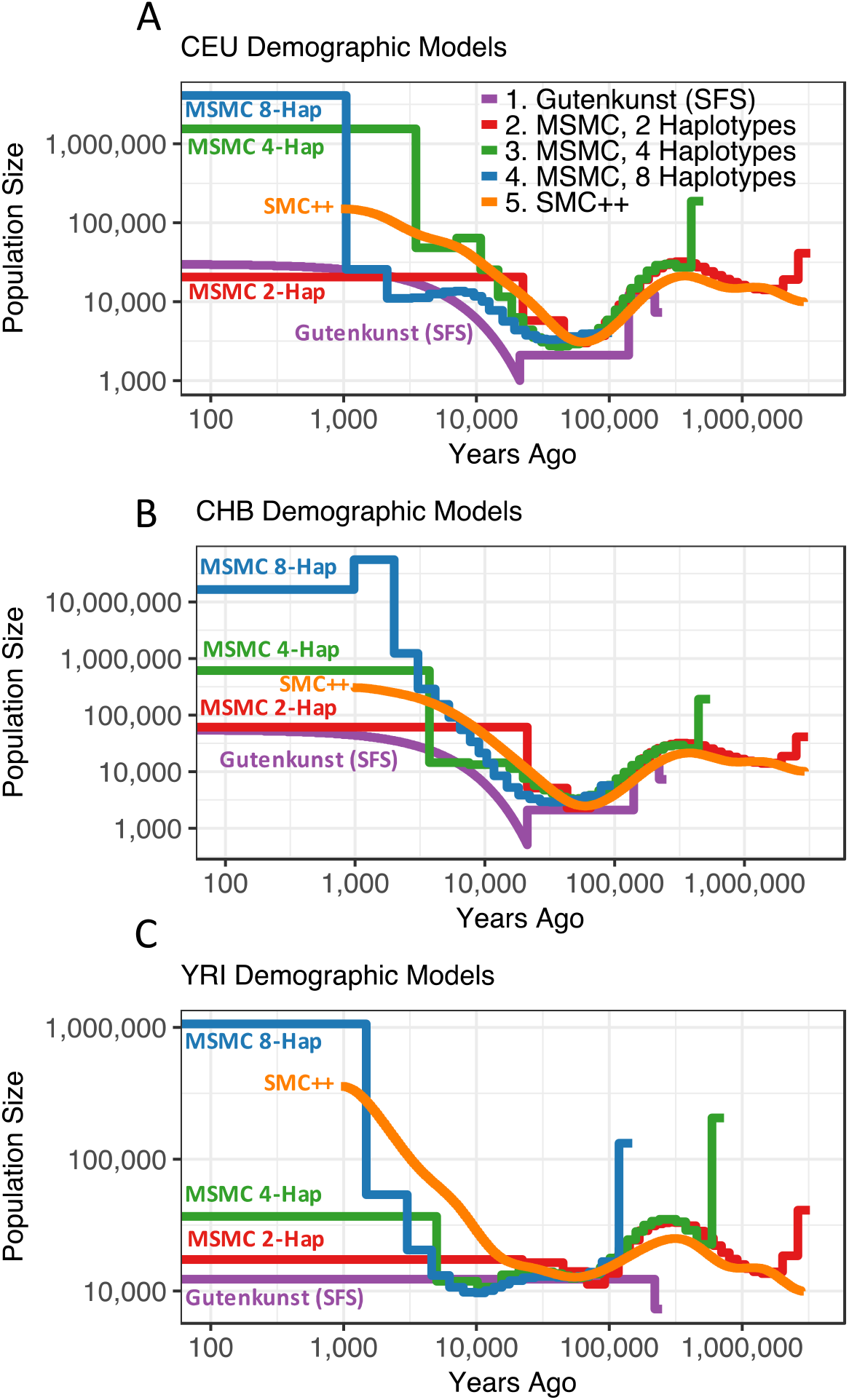
Demographic histories for the CEU (A), CHB (B), and YRI (C) populations. Trajectories are log-scaled and in terms of physical units (diploid individuals and years). Models were either inferred using SFS-based methods (“Gutenkunst”) by Gutenkunst *et al.* (2009), from a sequentially Markovian coalescent-based approach (“MSMC”) from two, four and eight haplotypes by Schiffels and Durbin (2014), or using a combined SFS and whole genome approach (“SMC++”) by Terhorst *et al.* (2017). The Gutenkunst models also include migration between all three populations, not depicted here. Models are scaled by the generation times used in each study (Gutenkunst *et al.* (2009): 25 years/generation; Schiffels and Durbin (2014): 30 years/generation; Terhorst *et al.* (2017): 29 years/generation).

Terhorst *et al.* (2017) analyzed the same populations with the combined whole genome and SFS method, SMC++, finding an ancestral size more in line with Gutenkunst *et al.*’s (2009) model, but with greater recent growth and ancestral bottlenecks more resembling the MSMC models (Figure 1). The reasons why these approaches to demographic inference yield different estimates remain poorly understood.

Here we leverage humans as a model system to perform an empirical comparison of the performance of whole genome, SFS, and combined methods of demographic inference. Specifically, we determine which published models of human demography described above (Figure 1) best fit the empirical distributions of genome-wide heterozygosity, LD decay, and the observed SFS.

We find that the models inferred using the SFS or the combined method SMC++ accurately recapitulate heterozygosity and the observed SFS. Among the MSMC models inferred by Schiffels and Durbin (2014), only the MSMC models based on a single genome were able to accurately recapitulate heterozygosity, and none of the MSMC models predicted an SFS that matched the empirical SFS. None of the demographic histories accurately predicted LD decay, but the histories derived from MSMC using four genomes (8 haplotypes), the SFS, and SMC++ based models fit better than the MSMC models based on one or two genomes. Our results provide a cautionary tale against the literal interpretation of demographic models inferred using one type of data, instead arguing for considering multiple summaries of the data when making detailed demographic inferences in non-model species.

## METHODS

### Published demographic models used in this study

We determined which, if any, of the published models of human demography (Figure 1) described below could accurately predict multiple summaries of the genetic variation data. Demographic models that fit the data well should produce patterns of genetic variation that match the empirical patterns in the data. We focused on three human populations: Utah residents with Northern and Western European ancestry from the Centre d’Etude du Polymorphism Humain (CEPH) collection (CEU), Han Chinese in Beijing, China (CHB), and Yoruba in Ibadan, Nigeria (YRI).

The first set of demographic models was jointly inferred for the three populations in ∂a∂i by Gutenkunst *et al.* (2009) using a three-population joint SFS based on data from intronic regions. Their model parameters were made available both in ∂a∂i and Hudson’s ms (Hudson 2002) format, and include gene flow between the three populations (here referred to as the “Gutenkunst” model).

The next nine models were inferred by Schiffels and Durbin (2014) using whole genome Complete Genomics (Drmanac *et al.* 2010) sequence data of two, four and eight statistically phased genomic haplotypes (1, 2 and 4 individual genomes) per population to infer demographic histories using MSMC (here referred to as the “MSMC 2-Haplotype”, “MSMC 4-Haplotype”, and “MSMC 8-Haplotype” models; **Supplementary Note 1**).

To analyze their models with ∂a∂i, we converted these nine demographic models (CEU, CHB, YRI populations, each based on two, four and eight haplotypes) into step-wise models of population size changes over small time intervals (**Supplementary Note 2, Figure S1**).

The final set of models was inferred by Terhorst *et al.* (2017) in SMC++, a combined SFS plus whole genome approach. For the whole genome portion of the analysis, they used high coverage sequence data from Complete Genomics, and generated an SFS based on a combination of 1000 Genomes and Complete Genomics whole genome data for each population (Drmanac *et al.* 2010; 1000 Genomes Project Consortium 2015; Terhorst *et al.* 2017). We converted these SMC++ models to ∂a∂i and ms format in the same manner as the MSMC models (here referred to as the “SMC++” models; **Supplementary Note 2**).

### Heterozygosity predicted by demographic models

We compared the distribution of expected heterozygosity from data simulated under each demographic model to empirical 1000 Genomes data from the same populations in order to determine which models most accurately predict this broad summary of the data (Figure 2; **Table S1**). While heterozygosity is a summary of the SFS, we considered it valuable to examine both statistics since information regarding the spatial correlation among SNPs along the genome is lost in the genome-wide SFS. The distribution of heterozygosity across windows of the genome retains some spatial information and is more similar to what is used by the MSMC inference approach.

**Figure 2.**
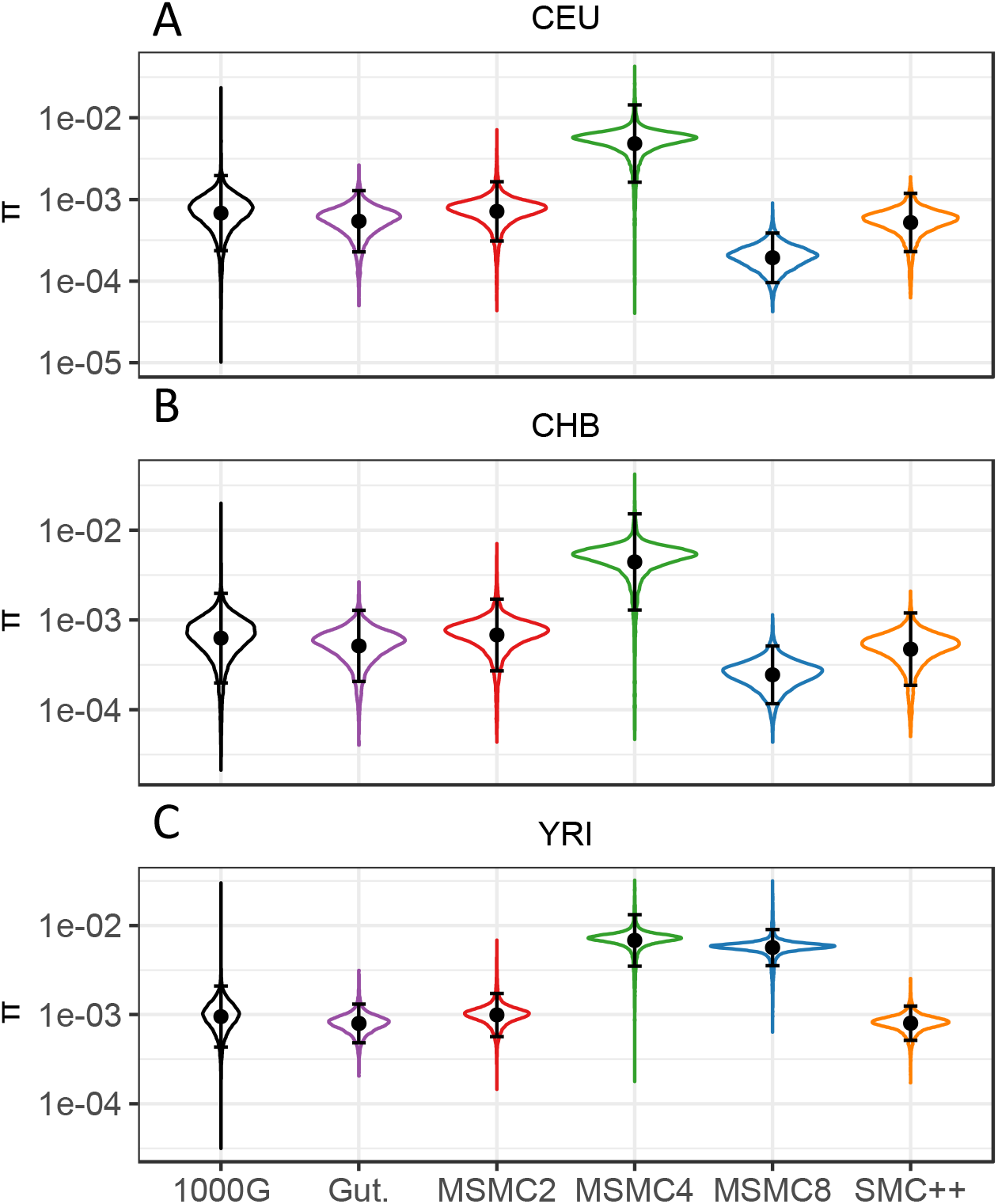
Kernel density distribution of expected heterozygosity (*π* per site). Heterozygosity was calculated across 100kb windows from whole genome 1000 Genomes project data for CEU (A), CHB (B), and YRI (C), and from 20,000 x 100kb blocks for data simulated under each demographic model. The black dot and bars indicate the mean ± two standard deviations for each distribution. Note the log-10 scaling on the y-axis.

#### Empirical heterozygosity

1000 Genomes data from the CEU, CHB and YRI populations were downloaded. Ten unrelated individuals per population (see **Supplementary Note 3** for sequence IDs) were randomly chosen so that comparisons could be made with Gutenkunst *et al.* (2009) empirical SFS based on 10 individuals, described below. For all our empirical analyses, only sites that passed the 1000 Genomes “Strict Mask” filter were considered (1000 Genomes Project Consortium 2015).

Expected heterozygosity per site (*π*) was calculated in non-overlapping 100kb windows from the whole genome data (**Supplementary Note 3**) as:

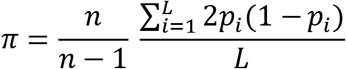

where *p* is the frequency of one allele, *L* is the total number of callable sites in the window, and *n* is number of sampled chromosomes (*n* = 20 for 10 diploid individuals).

Because genetic variation can be affected by linked natural selection (Gazave *et al.* 2014; Schrider *et al.* 2016), we also calculated expected heterozygosity for a set of 6333 x 10kb neutral windows that were selected using the Neutral Region Explorer (NRE) (Arbiza *et al.* 2012) (**Supplementary Note 3; Figure S2**). The NRE is a useful tool that allows for the quick identification of putatively neutral regions that have high recombination rates and high 5-values (indicating less linked selection). For the full set of parameters used in selection of putatively neutral regions, see **Supplementary Note 3**.

#### Simulated heterozygosity

For each demographic model, whole genome data for 10 individuals were simulated in MaCS (Chen *et al.* 2009) over 20,000 x 100kb independent blocks, each with a different recombination rate drawn from the distribution of recombination rates calculated by Phung *et al.* (2016) from the pedigree-based genetic map assembled by the deCODE project (Kong *et al.* 2010). Additionally, 6300 x 10kb independent blocks per 10 individuals were simulated for comparison to the neutral regions from the 1000 Genomes dataset (1000 Genomes Project Consortium 2015). Each 10kb block was simulated using a recombination rate matched to that of one of the empirical neutral 10kb windows, linearly interpolated from the deCODE project (Kong *et al.* 2010). For both sets of simulations, the expected heterozygosity across the 10 individuals was calculated using the equation above in msstats (Hudson 2002).

### Linkage disequilibrium decay predicted by demographic models

We calculated LD between pairs of SNPs using genotype data from 10 individuals from each of the four populations in the 1000 Genomes Project data. We removed singletons and sites where all ten individuals were homozygous for the reference allele and then calculated genotype r^2^ using vcftools (Danecek *et al.* 2011). All pairs of SNPs were then placed into bins based on their physical distance (bp) between each other, from 0-1000bp (bin 1) to 50,000-51,000bp (bin 51). Within each bin, the average *r*^2^ was calculated by dividing the sum of *r*^2^ values of each pair of SNPs in the bin by the total number of SNP pairs in that bin.

The same procedure was carried out for the data simulated in MaCS (Chen *et al.* 2009) that were used for the calculations of heterozygosity above. The MaCS output was converted to vcf format using a custom bash script. Genotype r^2^ was calculated in vcftools (Danecek *et al.* 2011) for each 100kb simulated window, the SNP pairs were binned by distance, and average r^2^ was calculated as described above. The MSMC 8-Haplotype YRI and MSMC 4-Haplotype CEU, CHB and YRI models have extremely large ancestral sizes, and so their simulations involve so many SNPs that the LD calculations become highly computationally intensive. Therefore, for these models only 5000 x 100kb blocks were used for LD decay calculations, with 20,000 x 100kb blocks used for the other models. We experimented with down-sampling the results and found no change in the LD decay curve due to the smaller amount of data.

To demonstrate that the use of the SMC’ approximation in the MaCS (Chen *et al.* 2009) simulator was not biasing our estimates of LD, we simulated data in the manner described above under a simple model of extreme population decline (from 100,000 ancestral individuals to 1000) using both MaCS and MSMS (Ewing and Hermisson 2010) (which does not use the SMC’ approximation) and ran it through the same LD decay pipeline used for our other simulated data (**Figure S3**).

### SFS predicted by demographic models

We used the diffusion approximation in ∂a∂i (Gutenkunst *et al.* 2009) to calculate the expected SFSs under the Gutenkunst, MSMC 2-Haplotype, MSMC 4-Haplotype, MSMC 8-Haplotype, and SMC++ models for the CEU, CHB and YRI populations. We compared the SFSs expected under each of these models both to the empirical SFS used by Gutenkunst *et al.* (2009) to infer the demographic histories of these three populations (“Observed (Gutenkunst)”, Figure 4–5) as well as to the SFSs based on low-coverage 1000 Genomes whole genome sequencing data (“1000 Genomes (Whole Genome)”, Figure 6) and SFSs based on putatively neutral regions in the 1000 Genomes dataset (“1000 Genomes (Neutral)”, Figure 6). We assessed the fit of different models to the observed SFS by comparing their log-likelihoods (see below, **Supplementary Note 4; Table 1, S2-S4**).

**Figure 3.**
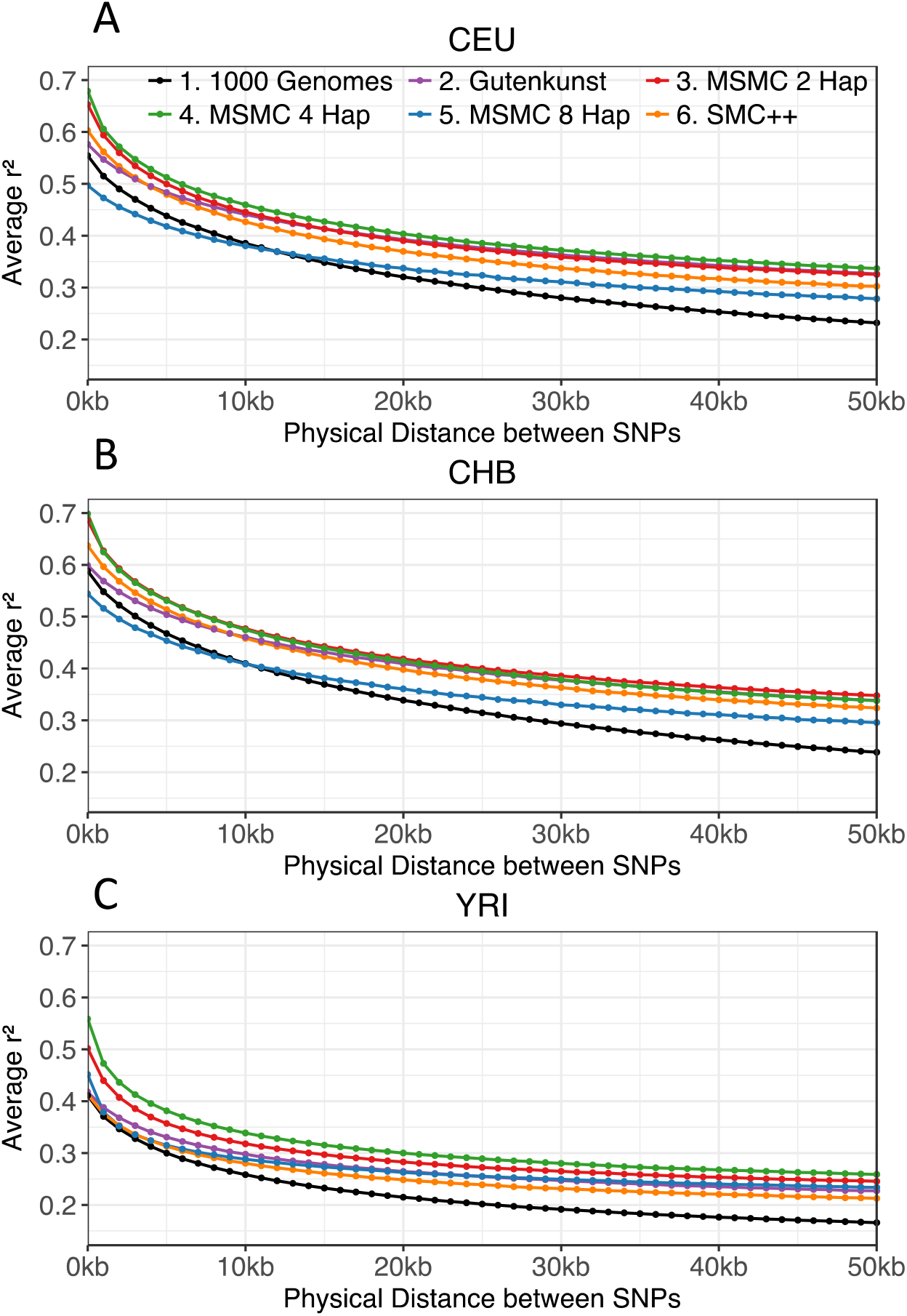
Linkage disequilibrium (LD) decay patterns. LD decay was calculated across 100kb windows from 1000 Genomes data and simulated data under each demographic model for CEU (A), CHB (B), and YRI (C). Pairs of SNPs are binned based on physical distance (bp) between them, up to 51kb. Average genotype *r*^2^ is calculated within each distance bin.

**Figure 4.**
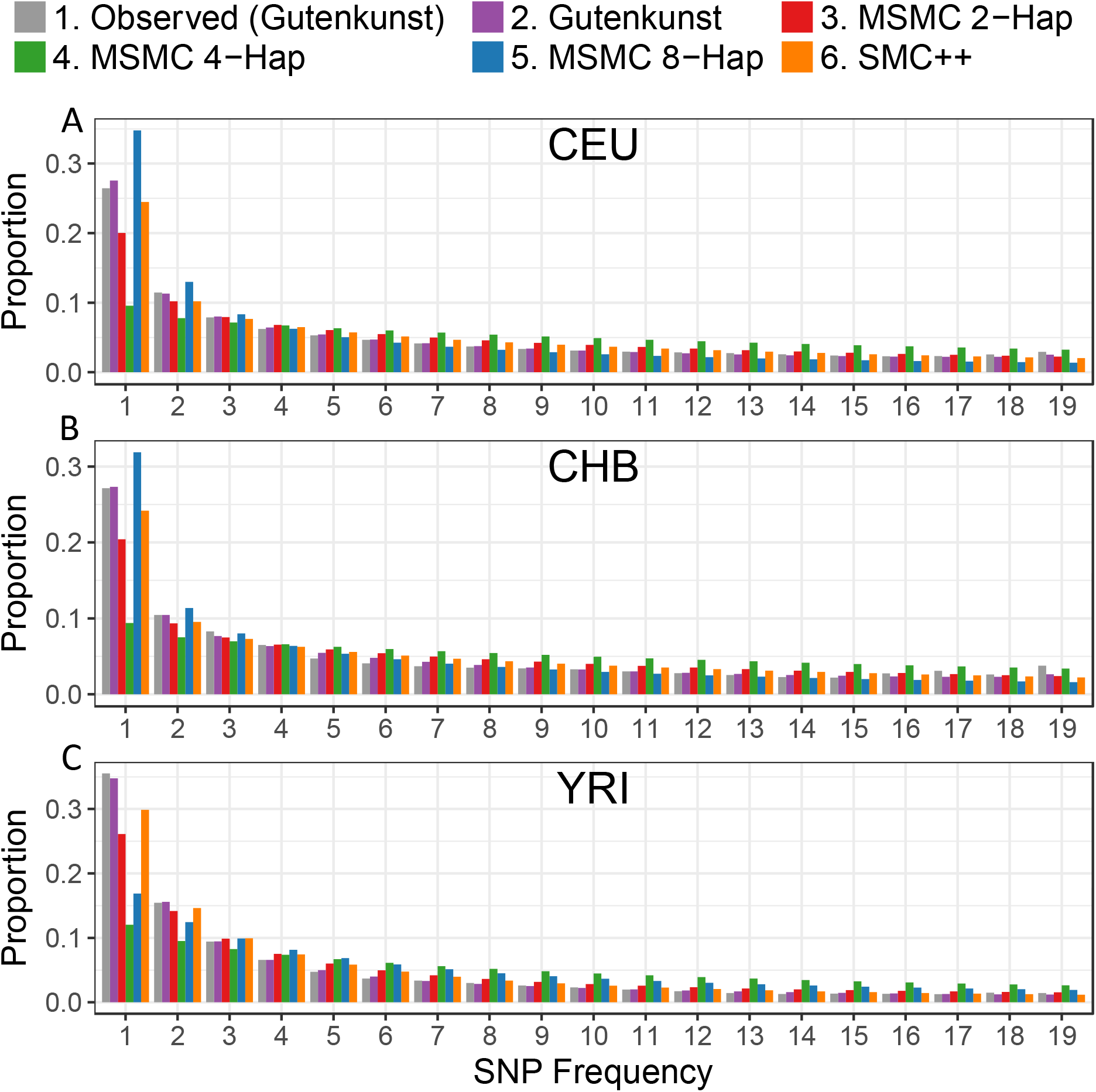
Unfolded proportional site frequency spectra for CEU (A), CHB (B), and YRI (C) populations. The “Observed” SFS is from noncoding sequence used by Gutenkunst *et al.* (2009) to infer demographic histories for these three populations. See **Figure S5** for scaling using alternative mutation rates.

**Figure 5.**
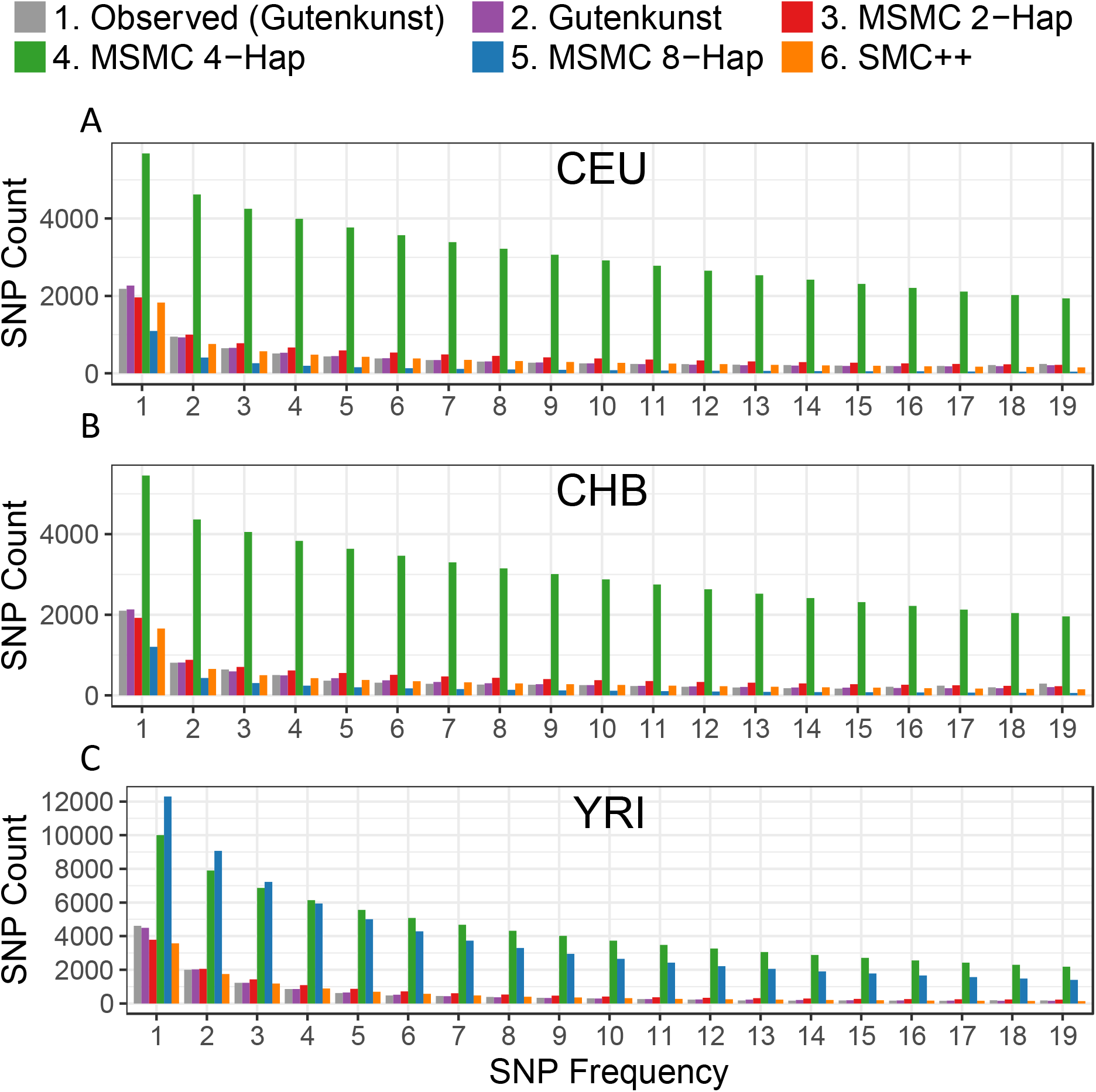
SNP count site frequency spectra using the counts of SNPs for the CEU (A), CHB (B), and YRI (C) populations. The “Observed” SFS is from noncoding sequence used by Gutenkunst *et al.* (2009) to infer demographic histories for these three populations. SFSs are scaled using the ancestral population size given by each model, the mutation rate used to scale each model by the authors and the sequence length of the empirical dataset (4.04Mb). See **Figure S6** for scaling using alternative mutation rates.

**Figure 6.**
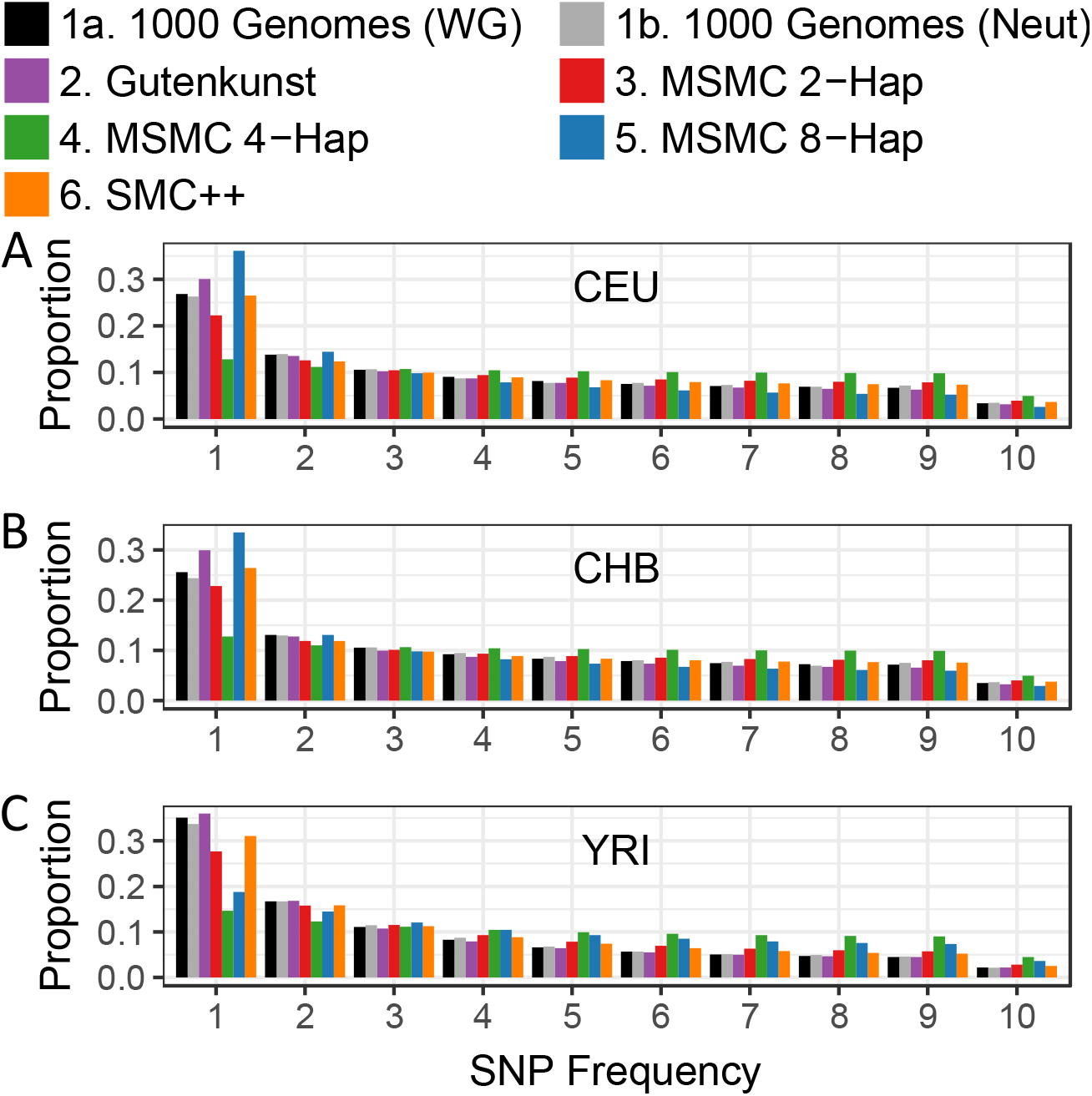
Folded proportional site frequency spectra for CEU (A), CHB (B), and YRI (C) populations. The “1000 Genomes (WG)” SFS is from low-coverage whole genome 1000 Genomes data, and the “1000 Genomes (Neut)” SFS is from 6333 x 10kb putatively neutral regions in the 1000 Genomes data.

**Table 1:** Multinomial log-likelihoods comparing the fit of various models to the observed SFS derived from Sanger sequencing data and used by Gutenkunst *et al.* (2009) for their inference (SFSs in Figure 4)

| CEU |  |  |
| --- | --- | --- |
| Model | Multinomial LL | $\Delta$ LL (Model - Data) |
| Data to Data<sup>a</sup> | -21546 | 0 |
| Gutenkunst<sup>b</sup> | -21555 | -9 |
| SMC++<sup>c</sup> | -21599 | -53 |
| MSMC 2-Hap<sup>d</sup> | -21698 | -152 |
| MSMC 8-Hap<sup>d</sup> | -21816 | -270 |
| MSMC 4-Hap<sup>d</sup> | -22760 | -1214 |
| CHB |  |  |
| Model | Multinomial LL | $\Delta$ LL (Model - Data) |
| Data to Data | -20154 | 0 |
| Gutenkunst | -20202 | -48 |
| SMC++ | -20277 | -123 |
| MSMC 8-Hap | -20343 | -188 |
| MSMC 2-Hap | -20370 | -216 |
| MSMC 4-Hap | -21411 | -1257 |
| YRI |  |  |
| Model | Multinomial LL | $\Delta$ LL (Model - Data) |
| Data to Data | -29630 | 0 |
| Gutenkunst | -29647 | -17 |
| SMC++ | -29779 | -150 |
| MSMC 2-Hap | -30003 | -373 |
| MSMC 8-Hap | -31282 | -1652 |
| MSMC 4-Hap | -32976 | -3346 |

a Denotes the best log-likelihood possible when replacing the proportions predicted by the model with the observed proportions from the SFS used in G utenkunst et al.’s (2009) study (see S upplementary Note 4).

b Denotes the model inferred by Gutenkunst et al. (2009) fit to the observed SFS

c Denotes the model inferred by Terhorst et al. (2017) using a combined whole genome and SFS approach

d Denotes the demographic models inferred by Schiffels and Durbin (2014) using MS MC on 2, 4 and 8 haplotypes

#### Empirical SFSs

The primary empirical SFSs used in our comparisons were produced by Gutenkunst *et al.* (2009) and used to infer the joint demographic histories of CEU, CHB and YRI populations in their study (“Observed (Gutenkunst)”). As described in their supplementary information, the joint SFS represents 4.04Mb of Sanger sequencing data from 10 diploid individuals per population for a total of 17,446 segregating SNPs polarized against chimp, with a correction for ancestral misidentification applied. We marginalized the SFS using ∂a∂i (Gutenkunst *et al.* 2009), in order to have one SFS per population (Figure 4, 5).

In order to make sure our results were consistent with SFSs derived from other sequencing methodologies and different genomic regions, we also generated folded proportional genome-wide and neutral SFSs from the 1000 Genomes data described above (“1000 Genomes (WG)” and “1000 Genomes (Neutral)”) (1000 Genomes Project Consortium 2015) (**Supplementary Note 3; Figure 6, S7**).

#### Expected SFSs under published demographic models

Expected SFSs for a sample size of 10 diploid individuals were calculated in ∂a∂i (2009) for each of the published demographic models extrapolating calculations across three grid points (40, 50, 60) (Figure 4, 5). To test whether the effect of differences in mutation rate between the studies may be responsible for discrepancies, we also considered an alternative scaling of the MSMC models using a higher mutation rate (**Supplementary Note 5**).

We generated both the proportional (**Figure 4, S5**) and absolute (i.e. SFS based on SNP counts) SFSs (**Figure 5, S6**). The proportional SFS was calculated by dividing each bin of the SFS output by ∂a∂i by the sum of the bins. The absolute SFS was calculated by scaling the SFS output by ∂a∂i (which is relative to *θ* = 1) by:

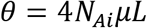

where *N_Ai_* is the oldest ancestral size inferred in each model and *L* is the sequence length (4.04Mbp), in Gutenkunst *et al.* (2009). *θ* for the Gutenkunst model used the authors’ preferred mutation rate, *μ* = 2.35x10^-8^ mutations per base per generation, and *θ* for the MSMC and SMC++ models used the authors’ preferred mutation rate of *μ* = 1.25x10^-8^ mutations per base per generation (see **Supplementary Note 5** for scaling using alternate mutation rates).

#### Assessing SFS fit

Log-likelihoods were calculated for each proportional SFS relative to the each of the three observed SFSs (Observed (Gutenkunst), 1000 Genomes (Whole Genome), and 1000 Genomes (Neutral)) using a multinomial log-likelihood (**Supplementary Note 4; Table 1, S2, S4**). The fit of different models was compared by examining their decrease in log-likelihood compared to that of each of the observed SFSs to itself (**Supplementary Note 4; Table 1, S2, S4**). Due to the uncertainty of singleton SNP calls using high-throughput sequencing data, log-likelihoods were calculated both with singletons and with the SFS renormalized without the singletons category when comparing to the 1000 Genomes SFSs (**Figure S7; Table S4**).

Log-likelihoods were calculated for each absolute SFS (in terms of SNP counts) using a Poisson likelihood relative to the Observed (Gutenkunst) SFS (**Supplementary Note 4; Table S3**).

### Effect of Uncertainty in Ancestral Population Size

To investigate whether changing the ancestral population size (*N_A_*) in the MSMC trajectories would result in SFSs that better fit the observed SFS, we adjusted the CEU MSMC 2-Haplotype model to have a variety of *N_A_* values. We also trimmed the model to remove ancient events (older than 225.5 kya) to better match the time period (in years) encompassed by the Gutenkunst *et al.*’s (2009) model. These adjusted stepwise models were then used to calculate the expected SFS in ∂a∂i, as above. **Supplementary Note 7** describes the values of *N_A_* used when testing the trimmed and untrimmed models (**Figure S10-S13**).

### MSMC Population Size Trajectories for Demographic Models Inferred from the SFS

To determine whether MSMC is capable of inferring a demography as complex as the one inferred in the Gutenkunst model, we used coalescent simulations to generate long chromosomal sequence data for each population under the Gutenkunst *et al.* (2009) inferred demographic model (see Gutenkunst *et al.*’s (2009) Figure 2B and Table 1 for full model), then ran MSMC on these simulated datasets to assess whether the program is capable of recovering the underlying demographic model.

Simulations were carried out using MaCS (Chen *et al.* 2009). For each population, we simulated 50 replicate “genomes,” made up of 80 independent 30Mb “chromosomes,” each made up of 300 linked 100kb recombination blocks, with per-block recombination rates calculated by Phung *et al.* (2016) from the pedigree-based genetic map assembled by the deCODE project (Kong *et al.* 2010).

Each simulated genome was then used for a separate MSMC inference, using the default parameters (Schiffels and Durbin 2014) (Figure 7A). To determine whether these inferred MSMC trajectories would lead to SFSs matching those predicted by Gutenkunst *et al.*’s (2009) model, the MSMC trajectories were averaged and the average was converted into a step-wise ∂a∂i model. This model was then used to calculate the expected SFS under the averaged model based on simulated data (Figure 7B-C). The multinomial and Poisson log-likelihoods for the proportional and SNP count SFSs were calculated as described in **Supplementary Note 4** (**Table S2, S3**).

**Figure 7.**
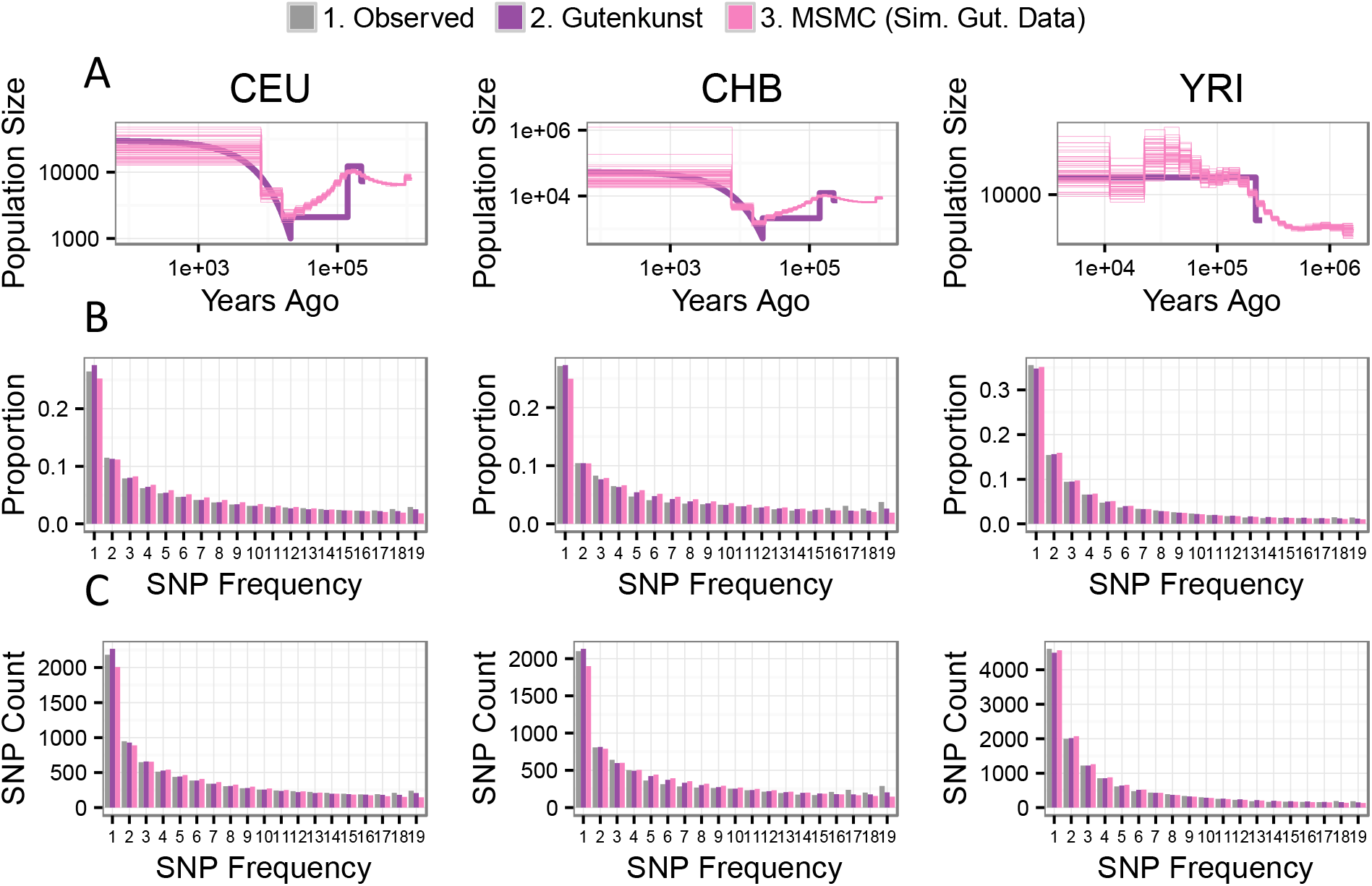
MSMC 2-Haplotype can accurately infer the demographic model predicted by Gutenkunst *et al.* (2009). (**A**) shows the results of running MSMC 2-Haplotype on 50 independent 2-haplotype datasets simulated under the Gutenkunst *et al.* (2009) model of human demographic history (“Gutenkunst,” heavy purple line). The resulting MSMC 2-Haplotype trajectories (“MSMC Sim. Gut. Data,” fine pink lines) show the MSMC trajectories inferred from these 50 datasets. Note that these trajectories accurately track the demographic model used to simulate the data. (**B**) and (**C**) show proportional and SNP count site frequency spectra for each population, respectively. The gray bars (Observed) denote the empirical SFS used by Gutenkunst *et al.* (2009). The purple bars denote the expected SFS under the inferred Gutenkunst demographic models. The pink bars denote the expected SFS under the average of the 50 MSMC 2-Haplotype demographic model trajectories for each population. Note that these three SFSs agree.

#### Extreme Recent Growth and Neanderthal Admixture

We simulated data under more complex demographic histories, first to explore MSMC 2-Haplotype and 8-Haplotype’s relative abilities to infer extreme recent growth, then to determine whether the addition of Neanderthal admixture may lead to MSMC trajectories resembling those inferred from real data by Schiffels and Durbin (2014) (**Supplementary Note 6; Figure S8-S9**).

### Data Availability

All code to simulate data under each demographic model and calculate heterozygosity and generate the SFS from simulated and empirical data are available on GitHub: github.com/LohmuellerLab/Compare_Demographic_Models

## RESULTS

We compared published models of demography for three human populations (CEU, CHB, YRI) inferred using different methods for demographic inference: (1) using the SFS in ∂a∂i (“Gutenkunst”) (Gutenkunst *et al.* 2009); (2) using whole genomes in MSMC (“MSMC 2, 4, 8-Haplotype”) (Schiffels and Durbin 2014); (3) using a combined SFS plus whole genome approach in SMC++ (“SMC++”) (Terhorst *et al.* 2017). The evaluation of the MSMC models involves three models per population because Schiffels and Durbin’s (2014) inference was carried out using 2, 4, or 8 chromosomal haplotypes (from one, two and four individuals), sometimes resulting in fundamentally different demographic parameter estimates. We evaluated whether the method’s performance was improved using certain numbers of haplotypes.

### Heterozygosity predicted by demographic models

The distribution of expected heterozygosity across 100kb and 10kb blocks was calculated from data simulated under each published demographic model for each of the three populations and compared to empirical distributions of heterozygosity based on whole genome and putatively neutral sequence data from the 1000 Genomes project.

We find that the Gutenkunst demographic model inferred from the SFS, the MSMC 2-Haplotype model and the SMC++ model all yielded distributions of heterozygosity that resemble the empirical whole genome distribution of heterozygosity, with MSMC 2-Haplotype fitting the mean most closely (Figure 2). However, we found that the higher haplotype MSMC models (MSMC 4-Haplotype and 8-Haplotype) yielded distributions of heterozygosity that were highly divergent from the empirical distribution (Figure 2; Table S1).

The MSMC 4-Haplotype models fit worst due to their extremely high inferred ancestral size across all three populations (Figure 1; **Table S2**; CEU: 187,514; CHB: 191,238; YRI: 205,845 individuals, compared to 4,000-40,000 individuals in the other models), with mean whole genome heterozygosity distributions nearly 7x larger than that of the empirical whole genome distribution (Figure 2; **Table S1**). The MSMC 8-Haplotye model for YRI infers a similarly large ancestral size and has a similarly high mean heterozygosity to the 4-Haplotype YRI model. The MSMC 8-Haplotype models for CEU and CHB, however, infer much lower ancestral sizes (CEU: 2,147, CHB: 5,666) (Figure 1). Due to the low ancestral size, these models also do not fit the empirical distribution well, yielding distributions of heterozygosity with means that are 2-4x lower than the empirical distributions.

When examining the 1000 Genomes data, we found that heterozygosity in the neutral regions was higher than that seen for the genome wide distribution of heterozygosity calculated in 10kb windows (**Table S1**; e.g. CEU mean heterozygosity per site, whole genome: 7.8x10^-4^ vs. neutral: 9.4x10^-4^), suggesting that natural selection has directly and/or indirectly affected genome-wide patterns of heterozygosity. When the published demographic models were compared to the neutral heterozygosity distributions, we found similar trends to those seen for the whole genome data (**Figure S2**).

### Linkage disequilibrium predicted by demographic models

None of the published demographic models could perfectly recapitulate the empirical LD decay curve (Figure 3). For SNP pairs less than 10kb apart, the MSMC-8 Haplotype model comes closest to the empirical curve for the CEU and CHB populations (Figures 3A and 3B), but underestimates the amount of LD, while all other models predict too much LD. The Gutenkunst and SMC++ models predict similar LD curves and are closer to the empirical curve than the MSMC 2-Haplotype and 4-Haplotype models. For YRI SNP pairs less than 10kb apart, SMC++ and MSMC 8-Haplotype predict similar LD decay curves and are close to the empirical distribution, with Gutenkunst still fitting better than MSMC 2-Haplotype and 4-Haplotype (Figure 3C). At distances greater than 10kb apart, all demographic models predict there to be more LD than seen in the empirical data (Figure 3).

We found that the lack of fit is not due to the use of the SMC’ approximation in the simulator MaCS (Chen *et al.* 2009), as both MaCS and MSMS (Ewing and Hermisson 2010), a coalescent simulator which does not use the SMC’ approximation, yielded highly similar LD decay curves when simulating data under the same simple population contraction model (**Figure S3**).

### SFS predicted by demographic models

Lastly, we examined which of the demographic models could match the SFS of the empirical data. To account for the possibility of overfitting the SFS-based Gutenkunst model to the SFS it was inferred from, we also compared all models to empirical SFSs based on low-coverage high-throughput 1000 Genomes sequence data from the same three populations.

#### Comparing to the observed Gutenkunst SFS

For each population, the SFSs predicted by the three MSMC models do not match the empirical proportional SFS from Gutenkunst *et al.* (2009), regardless of the mutation rate or number of genomes used (**Figure 4, S5; Table 1, S2**). The expected SFS based on the Gutenkunst *et al.* (2009) demographic history matches the observed SFS closely, being only 9 log-likelihood units worse than the best possible fit (comparing the empirical SFS to itself) for CEU, 48 units worse for CHB, and 17 units worse for YRI (Table 1). In comparison, the best fitting MSMC models for each population are 152, 188 and 373 log-likelihood units below the best possible fit (Table 1). The combined whole genome plus SFS method SMC++ has an intermediate fit, with a log-likelihood well below the Gutenkunst model, but consistently better than any of the MSMC models (Table 1).

Interestingly, there is not consistent improvement in fit to the observed SFS when increasing the number of individuals used for the MSMC inference. For each population, the 4-Haplotype model has the worst fit (Figure 4; Table 1). For CEU and YRI, the MSMC 2-Haplotype models fit best of the MSMC models, but both are over 100 log-likelihood units worse than the Gutenkunst model. For CHB, the 8-Haplotype model fits best, but is still 140 units worse than the Gutenkunst model (Table 1).

The above comparisons considered the proportions of SNPs at specific frequencies in the sample. We also performed a comparison of the number of SNPs in each bin of the SFS, the absolute SFS, to the observed absolute SFS used in Gutenkunst *et al.*’s (2009) inference using a Poisson likelihood. The absolute SFS expected under the Gutenkunst *et al.* (2009) model fits the observed SFS best (Figure 5; Table S3), and is only 9, 49 and 17 log-likelihood units below the best possible fits for CEU, CHB and YRI models, respectively. The SMC++ models have the next best fit to the absolute SFS, but come 86 (CEU), 176 (CHB) and 193 (YRI) log-likelihood units below the best possible fit, followed by MSMC 2-Haplotype which fell 278 (CEU), 378 (CHB), and 455 (YRI) below the optimal fit (Table S3). In all three populations, the MSMC 4-Haplotype and 8-Haplotype models are thousands of log-likelihood units worse than the best possible fit, showing no improvement based on using a larger number of individuals in the inference (Table S3). The over-estimation of SNPs in the 4-Haplotype model is due to the model’s extremely high predicted ancestral size (around 200,000 individuals for each population) (Table S3).

For both the proportional and absolute SFSs, we found that rescaling the models using a higher mutation rate did not produce large qualitative differences in how the MSMC models fit the observed (Gutenkunst) SFS (**Supplementary Note 5; Figure S4-S6**).

#### Comparing to the folded low-coverage 1000 Genomes SFS

To avoid giving the Gutenkunst model an unfair advantage by fitting all models to the SFS used to infer that particular model, we also compared all models to proportional folded SFSs based on whole genome and neutral data from the 1000 Genomes project (Figure 6, **S7**). The fit to the empirical singletons bin was poor for all models, except for SMC++, which was, in part, fit to an SFS based on 1000 Genomes data. Calling singletons is notoriously difficult in low-coverage data, making that bin the least reliable in the 1000 Genomes data (Kim *et al.* 2011; Nielsen *et al.* 2011; Han *et al.* 2014; 2015). We therefore calculated likelihoods for all models relative to the data both with singletons included and again with the SFSs renormalized without the singletons category (**Figure S7; Table S4**).

For YRI, the Gutenkunst model is the best fitting model for the whole genome and neutral 1000 Genomes SFSs, both with and without singletons, with all other models having a much worse fit (the next best model, SMC++, is hundreds to thousands of log-likelihood units below the fit of the Gutenkunst model) (Figure 6C; **Table S4**). For CEU and CHB, if singletons are included, SMC++ fits the whole genome and neutral 1000 Genomes SFSs best. For CEU, the Gutenkunst model then fits second-best, with the MSMC models far behind (Figure 6A; **Table S4**). For CHB, the MSMC 2-Haplotype fits second-best after SMC++, with the Gutenkunst model coming third, but both are over 10,000 log-likelihood units below SMC++ (Figure 6B; **Table S4**). If singletons are excluded for CEU and CHB, then the Gutenkunst model fits best, with SMC++ coming in second, and the MSMC models all ranking far below (**Table S4**).

### Effect of Uncertain Ancestral Population Size

The accuracy of ancient ancestral population sizes, particularly more than 3 million years (>100,000 generations) ago, using the whole genome-based methods remains unclear (Li and Durbin 2011). As discussed above, the MSMC 2-Haplotype and 4-Haplotype models infer large ancestral sizes for each population that are not supported by previous inferences of human demographic history (Adams and Hudson 2004; Keinan *et al.* 2007; Boyko *et al.* 2008; Gutenkunst *et al.* 2009; Nielsen *et al.* 2009; Gravel *et al.* 2011). We hypothesized that these extreme ancestral sizes, as well as ancient bottlenecks and population growth (the signature “humps” of MSMC trajectories), which do not appear in demographic models inferred using other methods, could be artifacts that are causing the SFS predicted by these models to deviate from the true SFS.

To test this hypothesis, we took the best fitting of the MSMC models, the CEU 2-Haplotype model, and carried out a series of adjustment experiments to determine whether changes to the model could provide a better fit to the observed SFS. Without adjusting the time period encompassed by the model, we altered the ancestral population size to a variety of values including those inferred by Gutenkunst *et al.* (2009) (**Supplementary Note 7; Figure S10-S11**). We also truncated the MSMC trajectory to remove ancient events and better match the time period (in years) encompassed by the Gutenkunst *et al.* (2009) model. We again adjusted the ancestral population size to a variety of plausible values (**Supplementary Note 7; Figure S12-S13**).

We found trimming away the ancient (older than ~225k years ago) part of the demographic trajectory and lowering the ancestral population size to 10,000 – 12,300 (compared to 41,261 inferred initially) dramatically improved the fit of the proportional SFSs predicted under these adjusted models to the Observed (Gutenkunst) SFS (**Figure S12; Table S5**). The best-fit model with ancestral size (*N_A_*) equal to 12,300 was brought to within 38 log likelihood units of the best possible likelihood (**Figure S12D; Table S5**), only 29 units below the Gutenkunst model. When repeating this procedure using the SFS based on counts, the SFSs under these adjusted models showed a different pattern of improvement. Here the untrimmed models that did *not* have ancient events >225 kya trimmed away, but had a lowered ancestral population size of 7,300-12,300, showed the most improvement (**Figure S11-S12**). However, their fit was still more than 100 log-likelihood units worse than the Gutenkunst model (**Figure S12; Table S6**).

### MSMC Population Size Trajectories for Demographic Models Inferred from the SFS

Given that the SFSs predicted by the demographic models inferred using MSMC do not fit the observed SFS, we examined whether MSMC is capable of recovering a complex demography such as the one inferred by Gutenkunst *et al.* (2009) from a single simulated genome. We find that MSMC performs relatively well at inferring the underlying demography from the simulated data. Figure 7A shows the underlying Gutenkunst demographic model for each population (purple) (as in the other Gutenkunst model simulations, migration is included in the model, but is not depicted in our diagrams), with the results of 50 independent MSMC inferences on each 2-Haplotype simulated dataset coming close to the underlying demography. However, sharp bottlenecks are inferred as long population declines (as noted by Li and Durbin (2011) and Schiffels and Durbin (2014)). Additionally, we found evidence of MSMC detecting a false spurt of growth in the YRI population 1350 generations ago (Figure 7A). Both of these phenomena were also noted by Bunnefeld *et al.* (2015).

The SFSs predicted by the demographic models inferred using MSMC on the simulated data fit the SFS expected under the Gutenkunst model and the observed Gutenkunst SFSs better than the MSMC demographic models inferred by Schiffels and Durbin (2014) (Figure 7B-C). The proportional MSMC simulated data SFSs were only 40, 74 and 10 log-likelihood units below the Gutenkunst model SFS (**Table S2**), with the SFSs based on SNP counts showing a similar pattern (**Table S3**). Therefore, if the Gutenkunst model is the true demographic model for human history, MSMC accurately captures the population size changes and produces an appropriate SFS.

It is well established that 2-haplotype whole genome-based inference (PSMC, MSMC 2-Haplotype, also known as PSMC’) is not able to detect recent demographic events (Li and Durbin 2011; Schiffels and Durbin 2014). However, the ability to detect recent growth by using more than two haplotypes in the inference is cited as a feature of MSMC (Schiffels and Durbin 2014). We ran MSMC 2-Haplotype and 8-Haplotype on datasets simulated under the Gutenkunst model and a Gutenkunst model plus extreme recent growth (**Supplementary Note 6; Figure S8**). Unsurprisingly, MSMC 2-Haplotype was not able to detect extreme recent growth. Its estimates of current population size were fairly accurate for the original Gutenkunst model (Figure 7A), but the method dramatically underestimated the growth for data simulated under the Gutenkunst + Growth model (**Figure S8**). The results from 8-Haplotype MSMC inference were most surprising. We found that for both models, MSMC 8-Haplotype inferred extreme recent growth as many as four orders of magnitude beyond that in the underlying model, with a high degree of variance between replicates (**Figure S8**). Despite the high degree of variance, the average of the MSMC trajectories all showed a strong upward bias in estimates of the recent past (**Figure S8**). While the ability to detect recent growth is meant to be a feature of MSMC, our findings indicate that the magnitude of growth may not be estimated well.

We had hypothesized that Neanderthal admixture could cause deviation between the MSMC and Gutenkunst demographic models, but found that the addition of Neanderthal admixture to our Gutenkunst model simulations did not substantively change the MSMC trajectories or expected SFSs (**Supplementary Note 6; Figure S9; Table S2, S3**).

## DISCUSSION

We tested which published models of human demographic history, inferred using either whole genome sequence data, the SFS, or a combined approach, can recapitulate multiple summaries of human genetic variation data. We found that no model was able to recapitulate all summaries of the data, but some models still performed better than others. In particular, none of the models was able recapitulate LD decay, but the Gutenkunst SFS-based models and the combined whole genome and SFS-based SMC++ models were able to recapitulate empirical heterozygosity and the SFS. MSMC 2-Haplotype was able to recapitulate heterozygosity, but not the SFS, and MSMC 4-Haplotype and 8-Haplotype could fit neither heterozygosity nor the SFS, though MSMC 8-Haplotype did fit LD decay slightly better than the other models. These results highlight the uncertainties of demographic inference from one, or even two, types of data and the need to assess the fit of demographic models using multiple summaries of the data.

We found that the models based on MSMC inference from 4 or 8 haplotypes did not improve the fit of the expected SFS compared to that based on two haplotypes; in fact, in most cases the 4- and 8-Haplotype models fit much worse than the 2-Haplotype models. The 4-Haplotype models for CEU, CHB and YRI and the 8-Haplotype model for YRI appear to fit poorly due to their extremely high ancestral sizes and ancient humps of growth and decline (Figure 1). The expected SFSs under the 8-Haplotype models for CEU and CHB show a skew toward low-frequency variants that may be due to their low ancestral size followed by extreme recent growth (Figure 1). We find that MSMC 8-Haplotype vastly overestimates recent growth in simulated data, which may be contributing to the lack of fit to the SFS (**Figure S8**). This result is at odds with the findings of Schiffels and Durbin (2014), who suggested that using eight haplotypes instead of two should increase accuracy of population size inference in the recent past, though they also noted a bias toward smaller ancient population sizes when using an increased number of haplotypes. Changing the scaling of the mutation rate did not generally help the MSMC models to fit the expected SFS better (**Figure S4-S6**). It is worth noting that the model inferred in SMC++ used the same mutation rate as MSMC, yet fit the empirical SFSs much better (Figure 4–6; Table 1, **S2-S4**), indicating that mutation rate differences between the whole genome and SFS-based studies is not the source of the discrepancies.

We found that in addition to not fitting the empirical SFS, the MSMC 4-Haplotype and 8-Haplotype models did not predict the genome-wide distribution of heterozygosity (Figure 2) which may be surprising as the genome-wide distribution of heterozygosity is a major feature of the data used by MSMC. The reason for the lack of fit for these models appears to be the extremely high ancestral size inferred in the 4-Haplotype models for all three populations and in the 8-Haplotype YRI model, and the low ancestral size inferred in the 8-Haplotype Models for CEU and CHB (Figure 1).

Since the most ancient size in the MSMC trajectory will have a large influence on heterozygosity and the SFS and the most ancient bin of the MSMC trajectory may be unreliable (Li and Durbin 2011; Schiffels and Durbin 2014), we explored the effect of altering this ancient size and removing ancient growth events in the CEU MSMC 2-Haplotype model. We found that selective trimming could improve the fit to the SFS (**Figure S10-S13**). However, the final bin of the model cannot explain all of the lack of fit of the MSMC models to the data as the CEU and CHB MSMC 8-Haplotype trajectories do not show the extreme ancestral sizes in the last bin, yet these models also dramatically deviate from empirical heterozygosity and the SFS. In other words, simple exclusion of the final high ancestral size is not sufficient to improve model fit to other summaries of the data. Our trimming experiments were only made possible by the abundance of human sequence data and demographic models previously fit to the data. Since many MSMC trajectories are calculated for species for which there is no prior information about ancient demographic history, the “informed trimming” we carried out is not a practicable solution to improve the reliability of MSMC inference.

While our results indicate that features of MSMC trajectories, particularly ancient events, should be regarded with caution, we also found that MSMC 2-Haplotype is able to accurately recapitulate a complex demography (with the exception of steep drops in population size, extreme recent growth, and some false periods of growth) from simulated data, supporting the validity of the method, at least for use on simulated data (Figure 7). Migration between populations did not appear to cause deviations in MSMC trajectories from the underlying model (Figure 7), nor did a small degree of Neanderthal admixture (**Figure S9**), indicating that MSMC is robust to small amounts of gene flow. The fact that the 2-Haplotype model based on real data did not fit the observed SFS very well (Figure 4–6; Table 1, **S2-S4**) suggests that the true underlying pattern of human demography is more complex than either type of inference (∂a∂i or MSMC) is capturing, potentially revealing weaknesses in both methods.

Alternatively, if the Gutenkunst *et al.* (2009) demographic model is largely accurate, biases or other factors that exist in real data but not in simulated data may be affecting MSMC inference, resulting in the method failing to recover an underlying demography that matches Gutenkunst *et al.*’s (2009) model. For example, Song *et al.* (2016) found that statistical phasing could affect MSMC estimates of population split times, and Nadachowska-Brzyska *et al.* (2016) found that per-site sequencing depth, mean genome coverage and the amount of missing data led to differences in PSMC curve amplitudes, expansions and contractions, and the timing and values of *Ne.* They therefore recommended only using samples with a mean genome coverage of ≥18X and < 25% missing data, and employing a per-site sequencing depth filter of≥10 (Nadachowska-Brzyska *et al.* 2016). The Complete Genomics genomes used by Li & Durbin (2011) were > 40X coverage (Drmanac *et al.* 2010), indicating that lack of coverage is not responsible for their divergence from estimates based on the SFS. However, the standards suggested by Nadachowska-Brzyska (2016) may not always be attainable in *de novo* genome projects, and thus, data quality issues may affect non-model organism PSMC and MSMC inferences more acutely. Future work should also examine the impact of artifacts of genome assembly errors and structural variants on PSMC inference. For example, collapsing duplicate regions of the genome on top of each other could result in regions of the genome having excess heterozygosity, which could in turn affect inference of demography.

We found that no model was able to accurately recapitulate the empirical distribution of LD decay. The lack of fit of the SFS-based models is perhaps unsurprising, as Harris & Nielsen (Harris and Nielsen 2013) found that the Gutenkunst model cannot recapitulate empirical IBS distributions (a finer-scale summary of the data related to LD), and Garud et al. (Garud *et al.* 2015) found that they could not recover empirical LD patterns in *Drosophila*, despite matching the SFS, number of segregating sites (S) and number of pairwise differences (*π).* Garud et al. (Garud *et al.* 2015) suggested the lack of fit could either be due to linked positive selection or to an incompleteness of the demographic model, demonstrating how models that fit some summaries of the data may not recapitulate others. It is more surprising that the MSMC 2-Haplotype and 4-Haplotype models do not fit the data well, as the method uses LD information in its inference, though different summaries of LD may be affected by demography in distinct ways (Plagnol and Wall 2006). Other possible factors that could lead to the lack of fit of all models to empirical LD decay patterns include the absence of natural selection, gene conversion, and fine-scale recombination hotspots in our simulations (Ardlie *et al.* 2001; Frisse *et al.* 2001; Wall and Pritchard 2003). Further, if the true mutation rate is actually smaller than the relatively high value used by Gutenkunst et al. (μ = 2.35x10^-8^ mutations/bp/generation), then the population sizes would have to be larger than those estimated by Gutenkunst et al. (2009). Larger population sizes would yield larger values of the population scaled recombination rate (*ρ*) than what was used in our simulations under the Gutenkunst model. Larger values of *ρ* would then lead to a decrease in LD in the simulations, which might better match the empirical LD decay curves.

Natural selection may affect both SFS and whole genome based methods of demographic inference. Li and Durbin (Li and Durbin 2011) found that masking exonic sequence did not alter PSMC trajectories. However, Schrider *et al.* (2016) examined the impact of selective sweeps on demographic inference using the SFS in ∂a∂i, approximate Bayesian computation (ABC), and PSMC and found that all three methods were influenced to varying degrees and in slightly different directions by the presence of selective sweeps, with ∂a∂i the most robust to these effects. This is a concern for published human demographic models as Gutenkunst *et al.* (2009) used noncoding sequence from autosomal genes in their study, which may be subject to linked selection (Gazave *et al.* 2014; Schrider *et al.* 2016). Schiffels and Durbin (2014) used whole genome sequences that included genic and non-genic regions some of which are certainly under selection. Thus, the sensitivity of these methods to selection may partially explain why both perform well on simulated data without selection, yet have such divergent results when run on empirical data.

Our results have implications for understanding human demographic history. First, there has been controversy concerning the presence of ancient bottlenecks (>100kya) in human populations (Takahata *et al.* 1995; Harpending *et al.* 1998; Takahata and Satta 1998; Hawks *et al.* 2000; Garrigan and Hammer 2006; Fagundes *et al.* 2007; Scholz *et al.* 2007; Blum and Jakobsson 2011; Sjödin *et al.* 2012). The inferred “humps” in the ancient portions of MSMC plots (Figure 1) tended to lend support to these ancient population size changes that appeared to be absent from SFS demographic estimates. Our results suggest that if these ancient population size changes did indeed occur, the resulting SFS would appear very different from the SFSs seen in human populations (Figure 4–6, **S10-S13**). The fact that they are not seen in the observed SFS suggests that either the size changes did not occur, and the inferred size changes are artifacts, or instead, the true demography is more complex than currently modeled using either approach. Our conclusion of finding little evidence for the ancient population size changes is supported by the study of Sjödin *et al.* (2012). They employed an approximate Bayesian computation approach to directly test models with ancient population size changes in Africa and found little support for such ancient bottlenecks.

Deep ancestral structure has been put forward as explanation for the humps detected by the whole genome-based methods by the developers of PSMC and others (Li and Durbin 2011; Henn *et al.* 2012; Mazet, Rodriguez, and Chikhi 2015; Mazet, Rodriguez, Grusea, *et al.* 2015; Orozco-terWengel 2016). While Blum and Jakobssen (2011) used the TMRCA to postulate an ancient bottleneck 150-kya, they also were not able to a reject a model of ancestral structure. Strikingly, Mazet *et al.* (2015) were able to perfectly recapitulate the human PSMC humps without invoking a single size change in the population by simulating data from a highly structured ancestral population (10 sub-populations) and modulating the amount of gene flow between these populations. Therefore, the large ‘population size changes’ inferred in MSMC, which cause the models not to match the empirical SFS, may in fact be due to complex structure and large-scale changes in gene flow. This ancient structure may have a large effect on MSMC trajectories and LD patterns, but may not strongly influence the SFS (see Figure 7 in Lohmueller *et al.* (2009)), potentially resolving the discrepancy between the methods (Henn *et al.* 2012).

Our work provides a cautionary tale for understanding population history in non-model organisms. Our results argue against a literal interpretation of “humps” and other jumps in MSMC plots as reflecting population size changes. This problem is exacerbated for putative ancient size changes. Given the ever-increasing generation of genomic data from non-model taxa and the application of whole genome-based approaches to such data (Meyer *et al.* 2012; Groenen *et al.* 2012; Zhao *et al.* 2012; Albert *et al.* 2013; Ibarra-Laclette *et al.* 2013; Orlando *et al.* 2013; Prado-Martinez *et al.* 2013; Nadachowska-Brzyska *et al.* 2013; Bosse *et al.* 2014; Freedman *et al.* 2014; Prufer *et al.* 2014; Hung *et al.* 2014; Nadachowska-Brzyska *et al.* 2015; Palkopoulou *et al.* 2015; Holliday *et al.* 2016; Nadachowska-Brzyska *et al.* 2016; Wang *et al.* 2016), our findings are especially concerning. We recommend employing other model-based types of demographic inference leveraging either SFS-based or other summary statistics in an ABC framework to test whether important demographic features suggested by PSMC or MSMC plots can be recapitulated using other features in the data. We also recommend, as done in Freedman *et al.* (2014), Song *et al.* (2016) and Cahill *et al.* (2016) that the PSMC or MSMC plots and TMRCA estimates be used themselves as summary statistics for model comparison, rather than the actual population size estimates. In other words, more complex demographic models can be simulated and tested to see whether they recapitulate the observed whole genome-based trajectories. Of course, this approach will not be successful if the trajectories are strongly influenced by bioinformatics artifacts or other features not captured within the simulations, such as natural selection. For both PSMC/MSMC and SFS-based inference methods, we also recommend testing whether the estimated models can predict multiple features of the data. Specifically, researchers should check whether their inferred model can recapitulate the genome-wide distribution of heterozygosity. The genome-wide distribution of heterozygosity may be the most practical and useful statistic for studies of non-model organisms that only have a handful of genomes available to them. SMC++ and other new approaches that leverage multiple types of data (Bunnefeld *et al.* 2015; Boitard *et al.* 2016; Weissman and Hallatschek 2017) are promising alternatives, though our results indicate that SMC++ still cannot recapitulate all summaries of the data.

Testing more complex demographic scenarios using multiple summaries of the data may help to resolve uncertainties about our own species’ history and will improve demographic inference for non-model organisms. Incorporating the potential complexity of possible demographic histories to produce models that better recapitulate the data may in fact present the greatest challenge.

## ACKNOWLEDGEMENTS

We thank Ryan Gutenkunst for providing us with the SFS from his paper, and Yun Song for his published demographic models. We also thank Diego Ortega Del Vecchyo and Bernard Kim for advice, and Christian Huber, Robert K. Wayne and Emilia Huerta-Sanchez for helpful comments on the manuscript. Joshua Schraiber and several anonymous reviewers made many insightful comments that strengthened our manuscript. This work was supported by National Institutes of Health (NIH) grant R35GM119856 to KEL and NIH Training Grant in Genomic Analysis and Interpretation T32HG002536 and National Science Foundation Graduate Research Fellowship Program to ACB. TNP was supported by National Institute of Health, under Ruth L. Kirschestein National Research Service Award (T32-GM008185).

## AUTHOR CONTRIBUTIONS

KEL and ACB conceived the study. ACB carried out all analyses based on the demographic models, and TNP carried out all empirical analyses based on the 1000 Genomes data. ACB generated all figures. ACB, TNP and KEL all participated in manuscript preparation.

